# Structural insights into filament recognition by cellular actin markers

**DOI:** 10.1101/846337

**Authors:** Archana Kumari, Shubham Kesarwani, Manjunath G Javoor, Kutti R. Vinothkumar, Minhajuddin Sirajuddin

**Affiliations:** Center for Cardiovascular Biology and Diseases, Institute for Stem Cell Science and Regenerative Medicine, GKVK Campus, Bangalore – 560065, India; National Center for Biological Sciences, TIFR, GKVK Campus, Bangalore – 560065, India; Manipal Academy of Higher Education, Manipal, India

## Abstract

Cellular studies of filamentous actin (F-actin) processes commonly utilize fluorescent versions of toxins, peptides and proteins that bind actin. While the choice of these markers has been largely based on availability and ease, there is a severe dearth of structural data for an informed judgment in employing suitable F-actin markers for a particular requirement. Here we describe the electron cryomicroscopy structures of phalloidin, lifeAct and utrophin bound to F-actin, providing the first high-resolution structures and comparison of widely used actin markers and their influence towards F-actin. Our results show that phalloidin binding does not induce conformations and lifeAct specifically recognizes ADP-actin state, which can be used as a sensor for distinguishing different nucleotide states of F-actin. The utrophin structural model aided designing minimal utrophin, which can be utilized as F-actin marker. Together, our study provides a structural perspective, where the binding sites of utrophin and lifeAct overlap with majority of actin binding proteins. Further offering an invaluable resource for researchers in choosing appropriate actin markers and generating new marker variants.

## Introduction

The cytoskeleton protein actin, can flux between globular (G-actin) and filamentous (F-actin) form, which is coupled to various cellular function. The polymerization kinetics of actin is dictated by the intrinsic nucleotide hydrolysis kinetics, phosphate release and ATP turnover cycle (1,2). The presence and absence of γ-phosphate at the nucleotide binding pocket induces conformational changes that leads to opening and closing of DNase binding loop (D-loop) respectively (1,3,4). In cells, several actin regulators can create an array of F-actin structures along with the nucleotide turnover and kinetics(5). For example, stress fibers, cortical actin, lamellopodia and filopodia (6); all of them show differential actin dynamics and are linked to a specialized actin mediated cellular process. To better understand these processes researchers often use fluorescent probes that label actin and visualize them using various microscopy methods (7). Broadly these markers can be categorized into fluorescent-tagged actin, toxins, peptides and proteins with actin binding domains (ABDs).

Fluorescent protein variants tagged at the amino-terminus of the actin gene has been used to label actin (7,8). The advantages of this approach include measurement of actin turnover in cells and *in vivo* whole organism studies, especially the control of expression with conditional and tissue specific promoters. The major disadvantage is the bulkiness of fluorescent proteins, which has been shown to impede incorporation of tagged G-actin into with F-actin (9). To overcome this, fluorescent toxins that bind to F-actin such as phalloidin (10) and jasplakinolide (SiR actin) (11) have been employed, of which phalloidin is perhaps the most widely used. Both phalloidin and jasplakinolide stabilizes F-actin (12), and structural investigations suggest that they both bind to same actin-binding site. Of these toxins, the jasplakinolide bound F-actin structure has been shown to mimic ADP-Pi actin state i.e., an open D-loop conformation (4,13). A similar conclusion for phalloidin could not be derived because the phalloidin bound F-actin structures determined so far, either have myosin (14) or filamin (15) bound to the filament, both of which overlap with the D-loop region.

The other commonly used reagent for F-actin labelling is lifeAct, a 17 amino acid peptide derived from yeast actin binding protein (16). Since its inception, the application of lifeAct to mark actin in cells elicits polarized responses among investigators; largely because lifeAct is shown to interfere with actin dynamics (17) and it fails to label certain actin structures in cells (8,18–20). LifeAct is also known to bind both G-actin and F-actin, with higher affinity towards the former form of actin (Riedl *et al*, 2008). However, a detailed structural analysis of lifeAct and actin interaction is still lacking.

In addition to toxins and peptides, the alternate method of actin labelling includes calponin homology domains (CH) that binds to actin, also known as tandem ABDs. The tandem CH1 and CH2 ABDs of utrophin (UTRN-ABD or UTRN-261, amino acids 1-261) have been successfully employed in F-actin visualization (21). Biochemical, structural and cell biological studies have proposed that the tandem arrangement of CH domain is important for F-actin binding (22–25). The crystal structure and biochemical experiments carried out with peptide fragments of utrophin suggest that CH1 domain has two actin binding sites and the third actin binding site was proposed to be in CH2 domain (26,27). Although the CH domains have high similarity among them, the linker between CH domains is variable and it is unclear whether the tandem CH architecture is important for utrophin:actin interaction. This is supported by truncation studies, which show the CH1 domain has higher affinity similar to the UTRN-ABD and the CH2 domain is important for solubility (28). Earlier studies have attempted the EM and helical reconstruction of UTRN bound to F-actin, however the importance of tandem CH domains could not be delineated due to its low-resolution data (23,24). Additionally, a shorter version of utrophin, UTRN-230 (amino acids 1-230) has been shown to specifically label Golgi actin (8) and nuclei actin (29), further questioning the actin binding sites and requirement of the tandem CH domain for actin interaction.

It has been well-documented and acknowledged in the field that, no one fluorescent actin marker is superior and all of them have certain limitations (8,17,19). Therefore, the accepted notion is that the choice of actin markers in investigations needs to be thoroughly thought through (7,18). However, all the studies have been limited to cell biology investigation and there is no structural study that has compared them systematically. In order to address these structural gaps, we employed electron cryomicroscopy (cryoEM) and helical reconstruction methods to determine the structures of actin markers bound to F-actin. Here we describe phalloidin, lifeAct and utrophin bound F-actin structure, representing toxin, peptide and protein markers widely used in actin labelling. We have also systematically compared the binding interface of these markers. The structures presented here will aid researchers in employing appropriate F-actin cell markers in their investigations.

## Results

### Phalloidin bound actin mimics the actin-ADP state

Phalloidin and jasplakinolide bound F-actin structures show that both share the same binding site (4,12,14). Jasplakinolide binding to actin induces the ADP-Pi like actin conformation state with an open D-loop, a nucleotide sensing region of actin (4). The available phalloidin structures are in complex with actomyosin or actin/filamin, where both myosin and filamin binding overlaps with the D-loop (14,15). Therefore, we determined the structures of F-actin-ADP apo and phalloidin bound using cryoEM and helical reconstruction (Supplement Figure 1 & Table 1) (Methods). Both of these F-actin structures contain ADP in the actin nucleotide binding site and thus forms the basis for the comparison of phalloidin induced conformational changes (Supplement Figure 2).

**Table 1:**
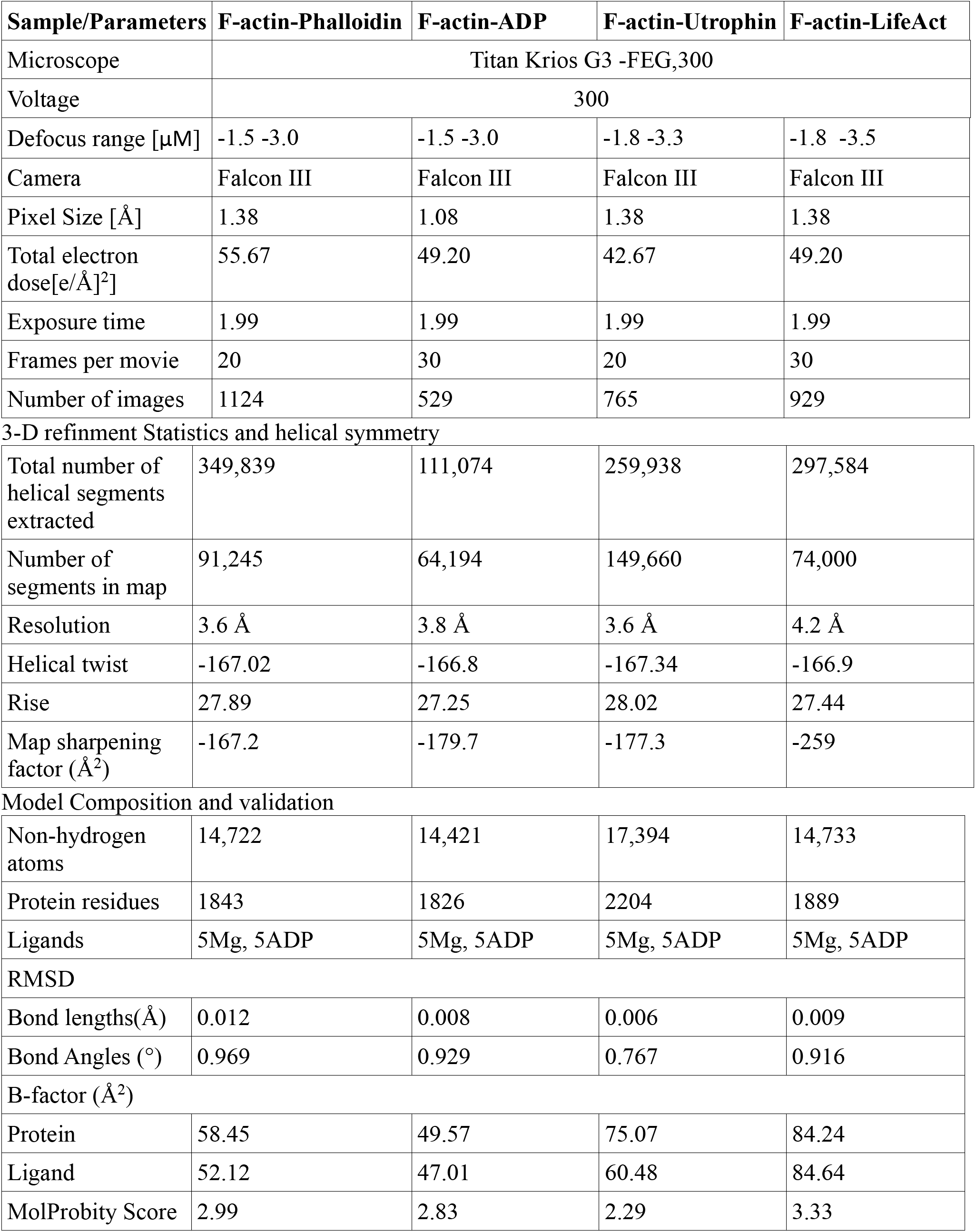
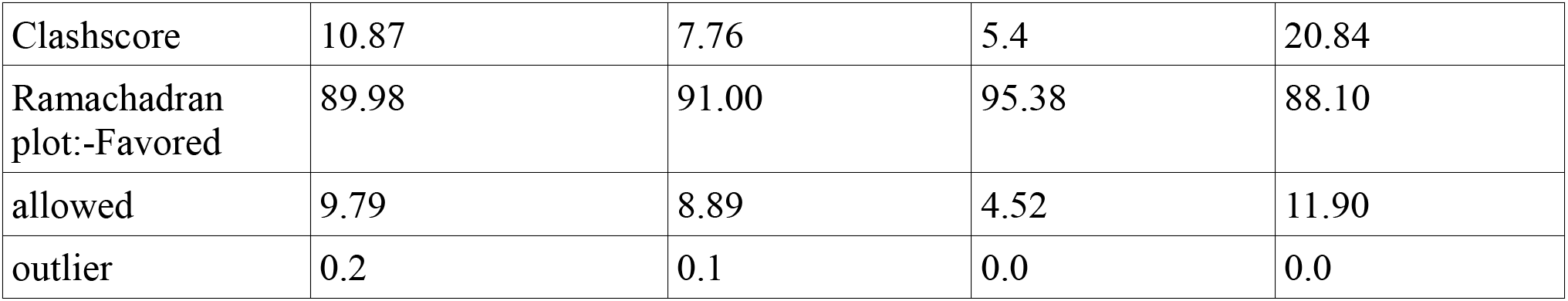

| Sample/Parameters | F-actin-Phalloidin | F-actin-ADP | F-actin-Utrophin | F-actin-LifeAct |
| --- | --- | --- | --- | --- |
| Microscope | Titan Krios G3 -FEG,300 |  |  |  |
| Voltage | 300 |  |  |  |
| Defocus range [$\mu\text{M}$] | -1.5 -3.0 | -1.5 -3.0 | -1.8 -3.3 | -1.8 -3.5 |
| Camera | Falcon III | Falcon III | Falcon III | Falcon III |
| Pixel Size [$\text{\AA}$] | 1.38 | 1.08 | 1.38 | 1.38 |
| Total electron dose [$\text{e}/\text{\AA}^2$] | 55.67 | 49.20 | 42.67 | 49.20 |
| Exposure time | 1.99 | 1.99 | 1.99 | 1.99 |
| Frames per movie | 20 | 30 | 20 | 30 |
| Number of images | 1124 | 529 | 765 | 929 |
| **3-D refinement Statistics and helical symmetry** |  |  |  |  |
| Total number of helical segments extracted | 349,839 | 111,074 | 259,938 | 297,584 |
| Number of segments in map | 91,245 | 64,194 | 149,660 | 74,000 |
| Resolution | 3.6 $\text{\AA}$ | 3.8 $\text{\AA}$ | 3.6 $\text{\AA}$ | 4.2 $\text{\AA}$ |
| Helical twist | -167.02 | -166.8 | -167.34 | -166.9 |
| Rise | 27.89 | 27.25 | 28.02 | 27.44 |
| Map sharpening factor ($\text{\AA}^2$) | -167.2 | -179.7 | -177.3 | -259 |
| **Model Composition and validation** |  |  |  |  |
| Non-hydrogen atoms | 14,722 | 14,421 | 17,394 | 14,733 |
| Protein residues | 1843 | 1826 | 2204 | 1889 |
| Ligands | 5Mg, 5ADP | 5Mg, 5ADP | 5Mg, 5ADP | 5Mg, 5ADP |
| RMSD |  |  |  |  |
| Bond lengths($\text{\AA}$) | 0.012 | 0.008 | 0.006 | 0.009 |
| Bond Angles ($^\circ$) | 0.969 | 0.929 | 0.767 | 0.916 |
| B-factor ($\text{\AA}^2$) | | | | |
| Protein | 58.45 | 49.57 | 75.07 | 84.24 |
| Ligand | 52.12 | 47.01 | 60.48 | 84.64 |
| MolProbity Score | 2.99 | 2.83 | 2.29 | 3.33 |
| Clashscore | 10.87 | 7.76 | 5.4 | 20.84 |
| Ramachadran<br>plot:-Favored | 89.98 | 91.00 | 95.38 | 88.10 |
| allowed | 9.79 | 8.89 | 4.52 | 11.90 |
| outlier | 0.2 | 0.1 | 0.0 | 0.0 |

The cyclic heptapeptide, phalloidin adopts a wedge like conformation and the binding pocket is buried in between three actin monomers (Figure 1A & Supplement Figure 2). Residues from the n+2^nd^ actin monomer was earlier reported to involve in hydrophobic contacts with phalloidin (14). However, a closer inspection of the binding site reveals that the nearest residue I287 from the third actin monomer (n+2^nd^ monomer) is at least 5Å away from phalloidin (Figure 1B & C). This indicates that phalloidin mainly interacts with two actin monomers and stabilizes the filament interface (Figure 1A & B). The binding pocket contains a mixture of hydrophobic and charged residues contributing to the phalloidin binding (Figure 1B and C). Phalloidin mainly interacts with E72, H73, I75, T77, L110, N111, P112, R177, D179 of n+1^st^ actin monomer and T194, G197, Y198, S199, F200, E205, L242 of n^th^ actin monomer. Superimposition of residues within the vicinity of phalloidin also does not show any major sidechain deviations between apo and phalloidin bound structures (Figure 1C).

**Figure 1:**
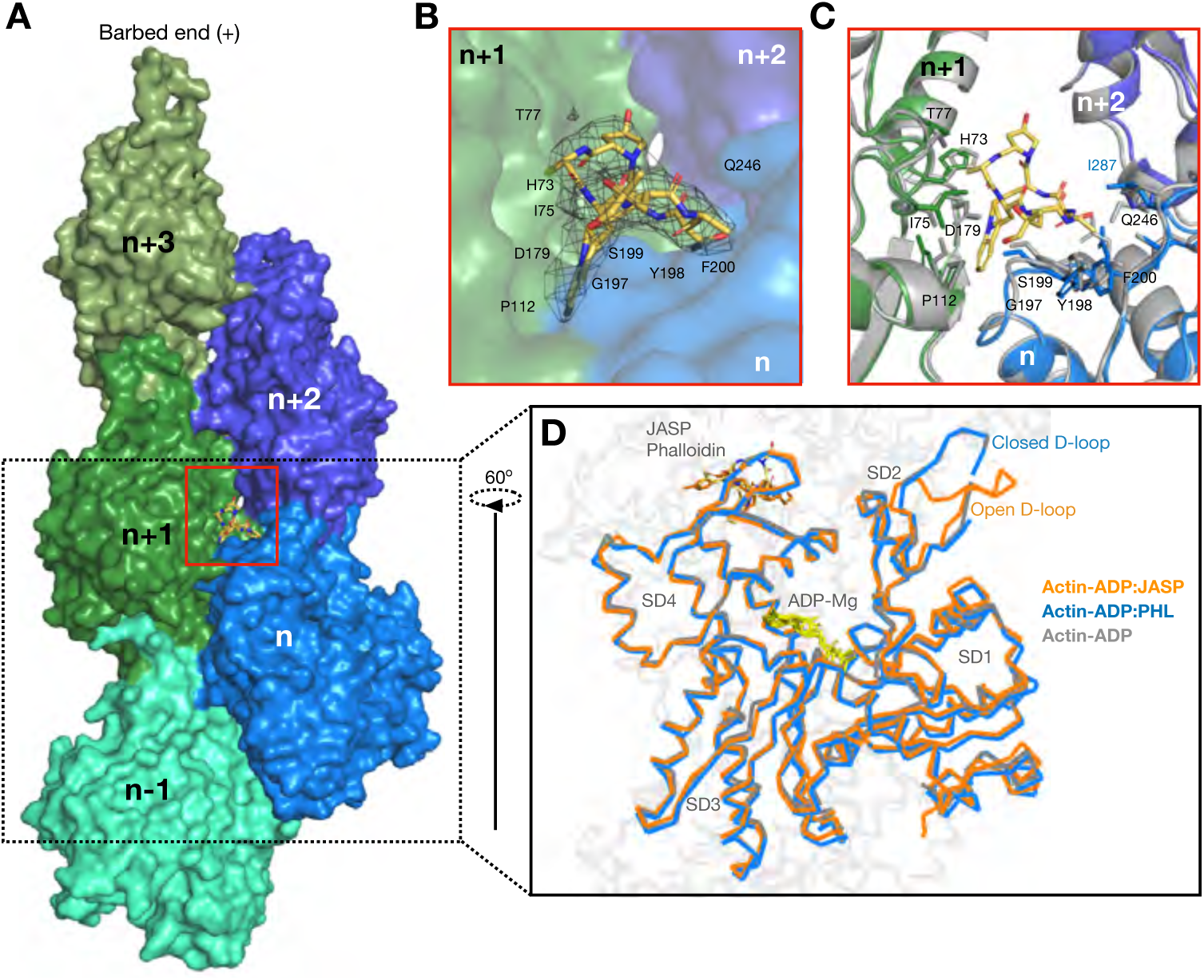
Phalloidin bound F-actin structure resembles ADP-actin state. **A.** Surface representation of F-actin ADP model, five monomers marked as n series from barbed to pointed end. A phalloidin molecule (yellow stick representation) bound between three actin monomers is highlighted. **B.** Closer view of phalloidin binding pocket as marked with red box in A. The density of phalloidin from EM map is shown around the ligand. **C.** Comparison of phalloidin binding pocket residues between Apo (in gray) and phalloidin bound (actin monomer colors as indicated in panel A) Key residues with their side chains and phalloidin are represented in stick representation. **D.** Overlay of F-actin ADP (gray), ADP/Phalloidin (blue) and ADP/Jasplakinolide (orange) shows the D-loop conformations across different structures as indicated.

Next, we superimposed and compared the apo, phalloidin and jasplakinolide bound F-actin-ADP structures (Figure 1D). Overall the architecture of the actin monomer does not deviate between the structures with the exception of the D-loop region (Figure 1D). In the ADP and ADP/phalloidin actin structures, the D-loop region remain in the closed state (rmsd 1.1 Å). While in in the ADP/jasplakinolide F-actin structure adopts an open conformation (4) (Figure 1D), the rmsd of D-loop between jasplakinolide versus phalloidin is 2.4Å. Because we have determined the undecorated F-actin structure, we conclude that phalloidin binding does not induce any conformational changes in actin and resembles the respective nucleotide state of F-actin (Figure 1D).

### LifeAct and F-actin interaction is mediated by hydrophobic contacts

Since its discovery, lifeAct has been widely used to detect actin using microscopy in cell biology studies (7). LifeAct is also known to influence actin dynamics and can bind both monomeric (G-actin) and F-actin (16). However, a detailed structural analysis of its interaction with actin is still lacking. To gain structural insights into the lifeAct:actin complex we determined a 4.2Å structure of lifeAct bound F-actin using cryoEM and helical reconstruction methods (Methods) (Supplement Figure 3 & Table 1). Similar to other actin markers, binding of lifeAct to F-actin does not alter the helical symmetry of the filament (Table 1).

LifeAct adopts a helical structure and binds stoichiometrically at the SD1 region of actin monomers and the carboxy-terminus of lifeAct extends towards the D-loop of the n-2^nd^ neighboring (pointed end) actin monomer (Figure 2A and B). The helical nature of lifeAct allows one to orient its hydrophobic residues, V3, L6, I7, F10 and I13 towards the actin (Figure 2B). On the actin front, a cohort of hydrophobic residues Y143, I345, L346, L349 and M355 mediates lifeAct binding (Figure 2B and 2C). An interesting feature is that the lifeAct binding pocket involves D-loop residues V45, M44 and M47 of the n-2^nd^ actin neighbor (Figure 2B and C). Together these residues form a hydrophobic pocket that can accommodate the phenyl sidechain group of the F10 lifeAct peptide (Figure 2B and 2C).

**Figure 2:**
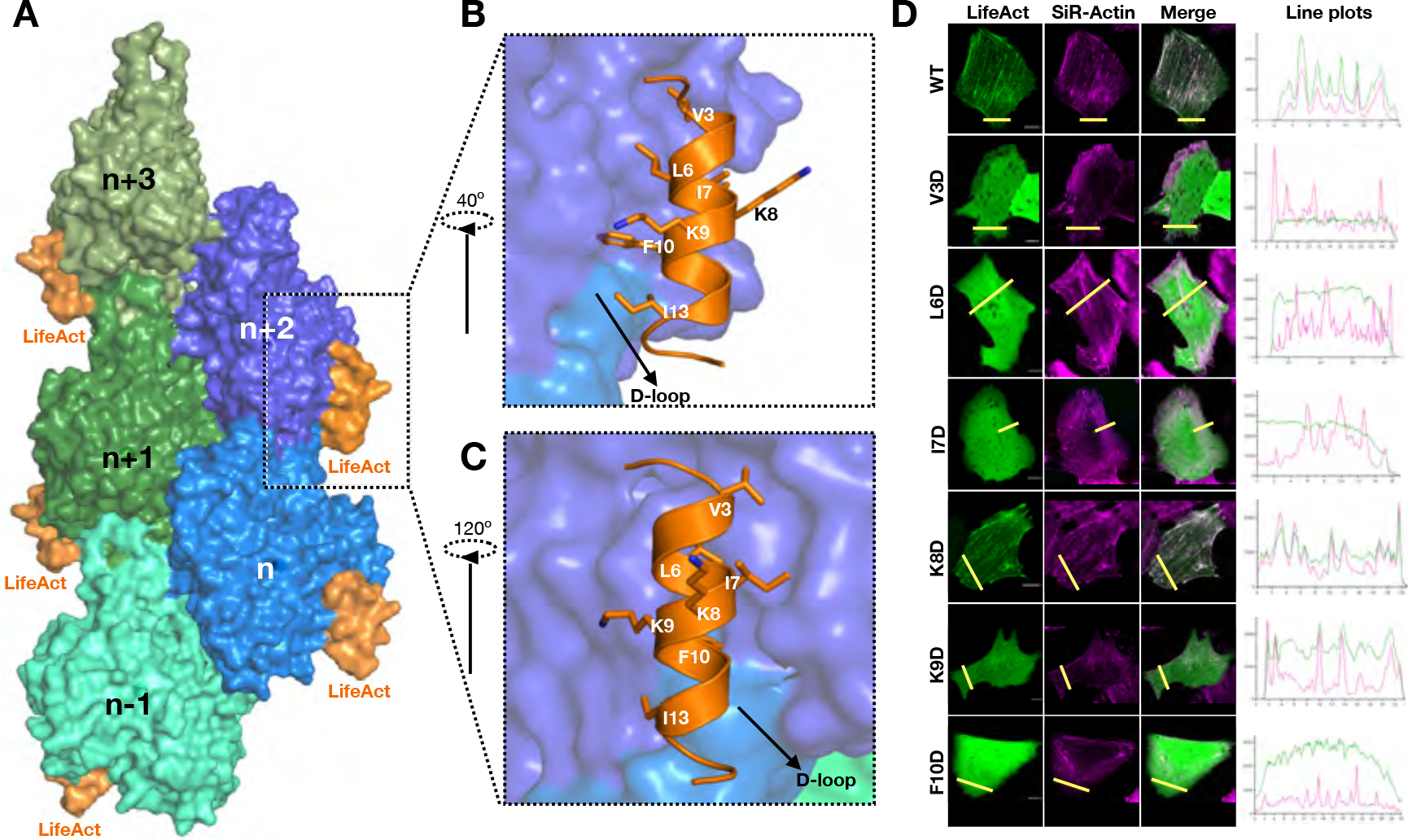
LifeAct structure, F-actin interaction and mutations. **A.** Surface representation of F-actin bound to lifeAct as indicated. **B & C.** Cartoon representation of lifeAct (orange) in closer look with n+1 and n+2^nd^ actin monomers (surface representation in blue), key interacting residues are highlighted. **C.** Schematic illustration of interaction between actin and lifeAct as indicated. **D.** Confocal images of U2OS cells transiently expressing lifeAct-GFP wild type and mutants of lifeAct residues interacting with F-actin, cells were additionally stained with SiR-actin to confirm the actin filaments. The line scan as indicated with yellow line on the cells shows the extent of lifeAct (green) and SiR-actin (magenta) co-staining of actin structures.

We then performed a mutagenesis experiment with lifeAct-GFP. The wildtype and mutant lifeAct were expressed in U2OS cells and their localization was imaged with actin structures (Methods). Here we chose V3, L6, I7, K8, K9 and F10 and replaced them with aspartic acid (Figure 2B and 2C). Colocalization with the SiR actin probe showed that only residues that mediate hydrophobic contacts with F-actin as described above drastically reduced binding to F-actin in cells (Figure 2D). Thus, further validating our structural observations of lifeAct and F-actin interaction.

### LifeAct senses the closed D-loop conformation

Since lifeAct peptide binding overlaps with the D-loop of the n-2^nd^ actin neighbor, we next probed the importance of D-loop conformation towards lifeAct and F-actin interaction. Comparison of open (jasplakinolide bound F-actin:ADP PDB: 5OOC) versus closed D-loop states (F-actin:ADP:lifeAct) (Figure 3A) suggests that the open D-loop state is incompatible for lifeAct binding (Figure 3B). Therefore, from the structural model of lifeAct:actin complex we reasoned that lifeAct may have preference towards different biochemical states of actin. We therefore prepared two batches of F-actin, one with phalloidin bound and the other with jasplakinolide bound, representing ADP and ADP-Pi actin state respectively (Methods) (Figure 1 D). The two distinct fluorescent F-actin populations were incubated together with varying concentrations of FAM-lifeAct peptide and visualized in the same reaction chamber using TIRF microscopy (Methods) (Figure 3C). At micromolar concentrations, we began to observe F-actin labelling by FAM-lifeAct, however the co-localization was favored towards the phalloidin F-actin form (Figure 3C and D). We quantified the fluorescence intensity ratio of FAM-lifeAct for phalloidin versus jasplakinolide F-actin (Methods) (Figure 3D), and found that a 3-4 fold fluorescence increase towards the phalloidin bound F-actin i.e., ADP actin. This trend and the fluorescent intensity ratio were observed at different concentrations of lifeAct (Figure 3C and D). The striking preference of phalloidin over jasplakinolide F-actin thus strongly suggests that lifeAct preferentially binds to the ADP state of F-actin.

**Figure 3:**
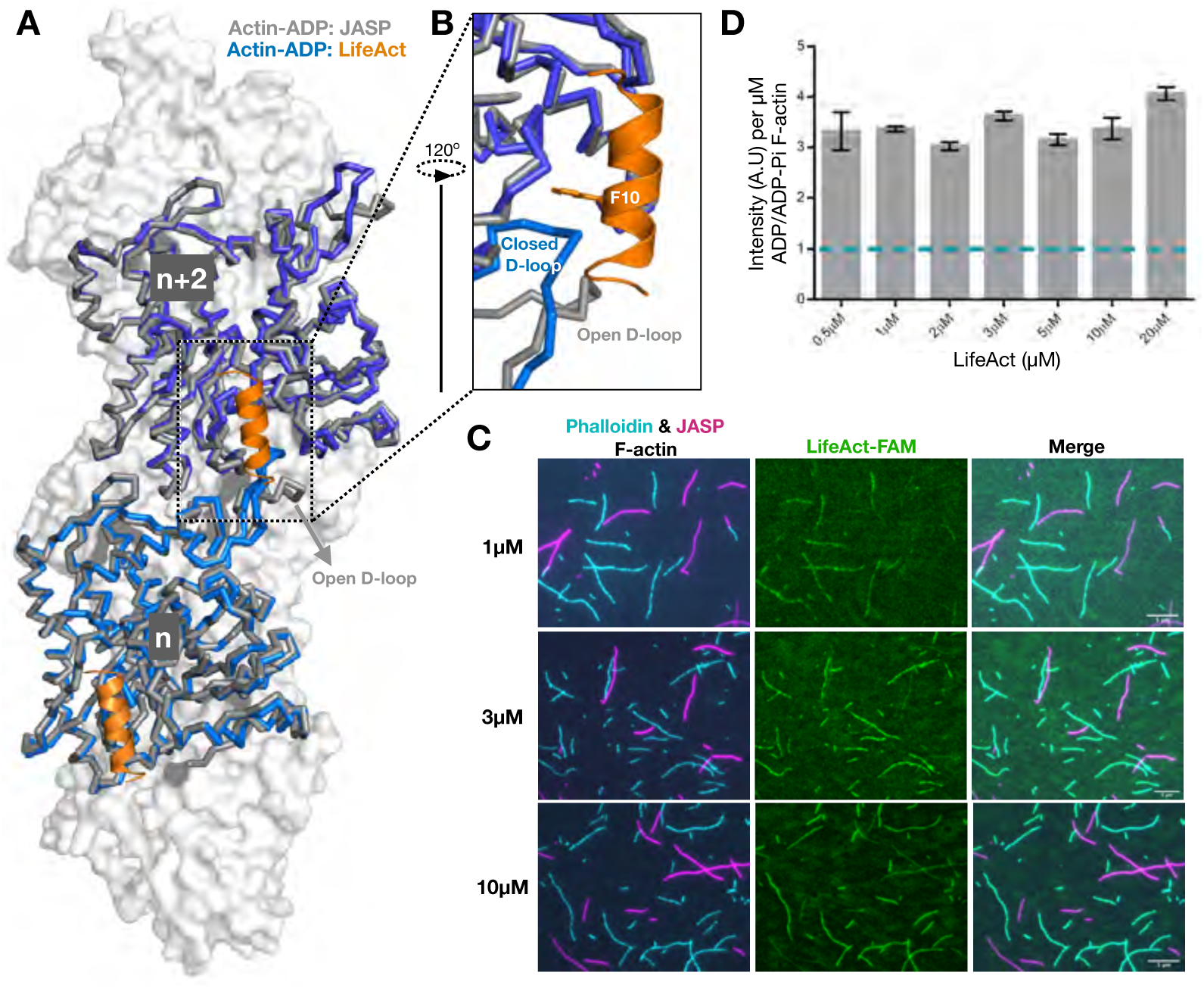
LifeAct recognizes closed D-loop state of F-actin. **A.** and **B.** Overlay of F-actin:ADP bound LifeAct (actin in blue and lifeAct in orange cartoon) and jasplakinolide (in gray). **C.** Representative TIRF images of lifeAct binding experiments as indicated. The remaining lifeAct concentration images are shown in Supplement Figure 4. **D.**Mean ratio of lifeAct fluorescent intensity bound to phalloidin and jasplakinolide F-actin (mean and s.e.m; n = 2 or 3 independent experiments with >50 actin filament for each set).

### The utrophin CH1 domain is sufficient for F-actin interaction

In our quest towards structural characterization of actin markers, we then focused on the utrophin actin binding domain, widely known as UTRN-ABD or UTRN261, amino acids 1 – 261. The UTRN-ABD contains two calponin homology domains (CH1 and CH2 domains) and previous structural and biochemical studies have proposed both the domains are necessary for actin interaction(22). Here we purified the UTRN-ABD:F-actin complex (Methods) and subjected it to cryoEM helical reconstruction methods and resolved to a structure to 3.6Å resolution (Methods) (Table 1 and Supplement Figure 4). The UTRN-ABD model was built from the available X-ray structure coordinates (PDB ID: 1QAG)(26) and the additional amino-terminal helix, which was partially disordered in the X-ray structure was built *de novo*. Although our cryoEM preparations contain complete UTRN-ABD protein (amino acids 1 – 261), in the final reconstructed map we could model only less than 50% of the utrophin, amino acids 18-135 corresponding to the CH1 domain (Figure 4A and Supplement Figure 4).

**Figure 4:**
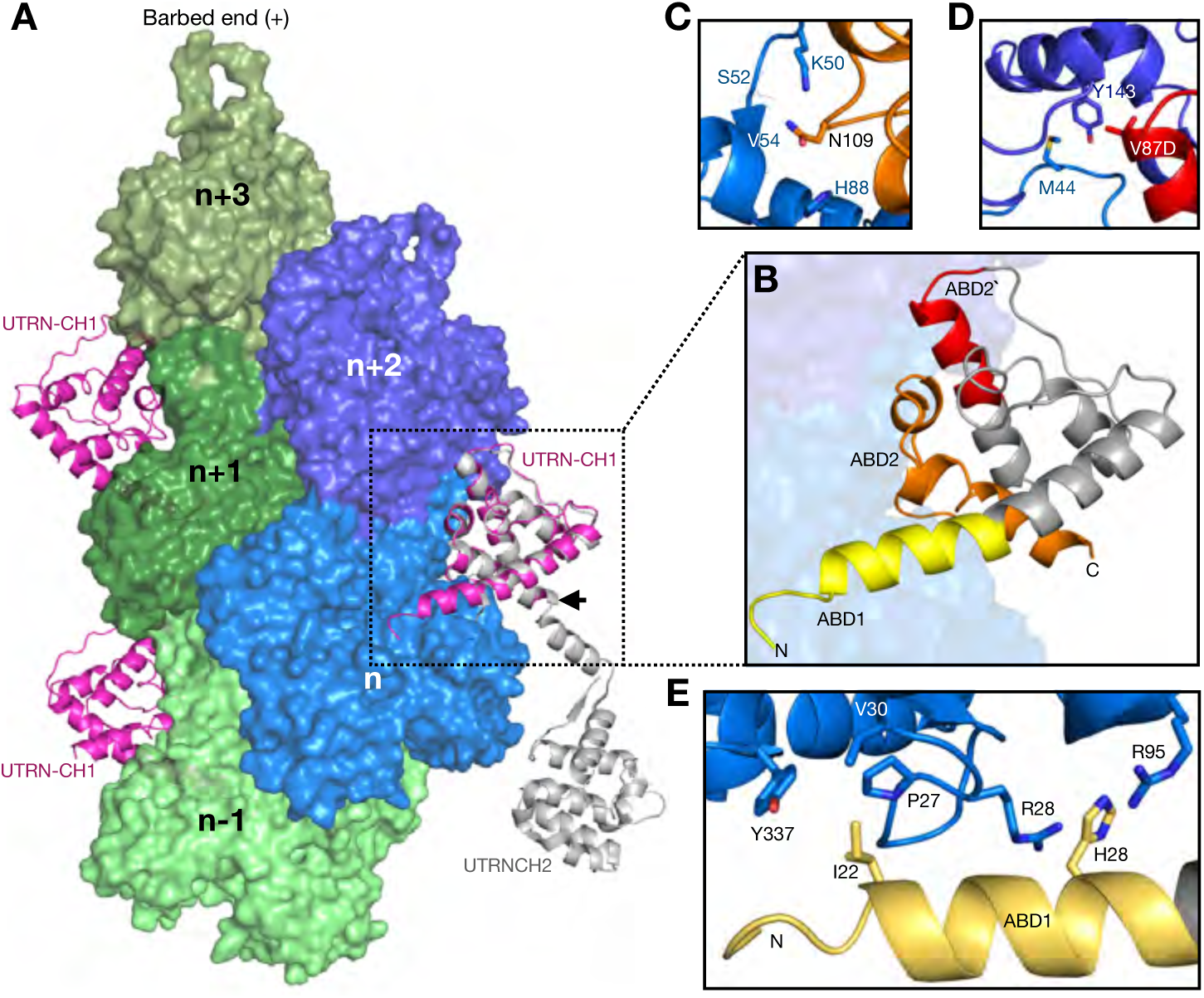
Utrophin CH1 domain structure and F-actin interaction sites. **A.** Surface representation of F-actin ADP, five monomers marked as n series from barbed to pointed end. The utrophin CH1 domain in magenta interacts with two adjacent actin monomers thus following the actin helical pattern. The crystal structure of dystrophin/utrophin in gray (1DXX) superimposed with cryoEM utrophin CH1 model, boundary of CH1 is marked by an arrow. **B.** Closer view of utrophin CH1 model, the yellow, orange and red region depicts ABD1, ABD2 and ABD2’ sites respectively. The ABD1 and ABD2 sites are restricted to n^th^ actin monomer, the ADB2’ site partially interacts with the neighboring n+2^nd^ actin monomer. **C, D & E.** Residual level information of key amino acids interacting with actin monomers from ABD1: I22, H29 (C); ABD2: N109 (D) and ABD2’ site: V87 (E).

Previous work subdivided the CH1 domain of UTRN-ABD into ABD1 (amino acids 31 – 44) and ABD2 (amino acids 105 – 132) (26). From our F-actin bound structure we could redefine the boundaries of ABD sites; ABD1 (amino acids 18-33), ABD2’ (amino acids 107-126) and a newly identified ABD site in between ABD1 and ABD2, named ABD2 (amino acids 84 – 94) (Figure 4B). ABD1 is the amino-terminal helix, which mainly interacts with the SD1 of the n^th^ actin monomer (Figure 4A & B). Both ABD2 and ABD2’ interacts with SD2, chiefly with the D-loop region of the n^th^ actin monomer, which remains in a closed conformation (Figure 4A & B). Additionally, ABD2’ is the only site that interacts with the SD1 of adjacent n+2^nd^ actin monomer (Figure 4A & B). The binding site and architecture of UTRN-ABD- CH1 is similar to the recently reported FLNa-ABD(15), however in our UTRN-ABD an additional amino-terminal helix is visible, extending towards the pointed end of the actin monomer (Figure 4A & B).

To validate our structural model of the UTRN-ABD:actin complex and the newly defined ABD1 (amino-terminal helix) and ABD2’ region we performed mutagenesis of key interacting residues (Figure 4C, D & E) and co-sedimentation assays with F-actin (Methods) (Supplement Figure 6). From the ABD1 helix, we chose residues that have their side chains facing towards actin; thus, I22 makes hydrophobic contacts with a cohort of P27, V30 and Y337 of actin residues, and H29 engages in a cation-pi interaction with R28 and R95 of actin. (Figure 4E). At the core of the CH1 domain (ABD2), we included N109, which makes electrostatic interactions with actin K50 and H88 side chains and the main chain carbonyl group of S52 and V54 (Figure 4C). Additionally, we included V87D from the newly identified ABD2’, which makes hydrophobic contacts with Y143 of the SD1 of the adjacent n+2^nd^ actin monomer and M44 from the D-loop of the n^th^ monomer (Figure 4D). In summary, we tested the following mutants, I22D and H29A from the ABD1 site, V87D and N109A for ABD2’ and ABD2 sites respectively (Figure 4B, C, D & E).

Co-sedimentation assays of mutants compared to the wildtype UTRN-ABD protein show more than 50 and 100 times decrease in binding constants for V87D and N109A mutants respectively (Figure 5A & B). The decrease in affinity by V87D and N109A mutants indicate that the core binding is mediated by the ABD2’ and ABD2 sites. Moreover, the reduced binding constants of the V87D mutant data indicates that the UTRN-ABD interacts with two neighboring (n^th^ and n+2^nd^) actin monomers and thus has the ability to bind to F-actin, but not actin monomers (22). Our mutation analysis also shows that the I22D, but not H29A has a profound impact in binding affinities, suggesting that the ABD1 (amino-terminal helix) might play an important role in actin binding (Figure 5B).

**Figure 5:**
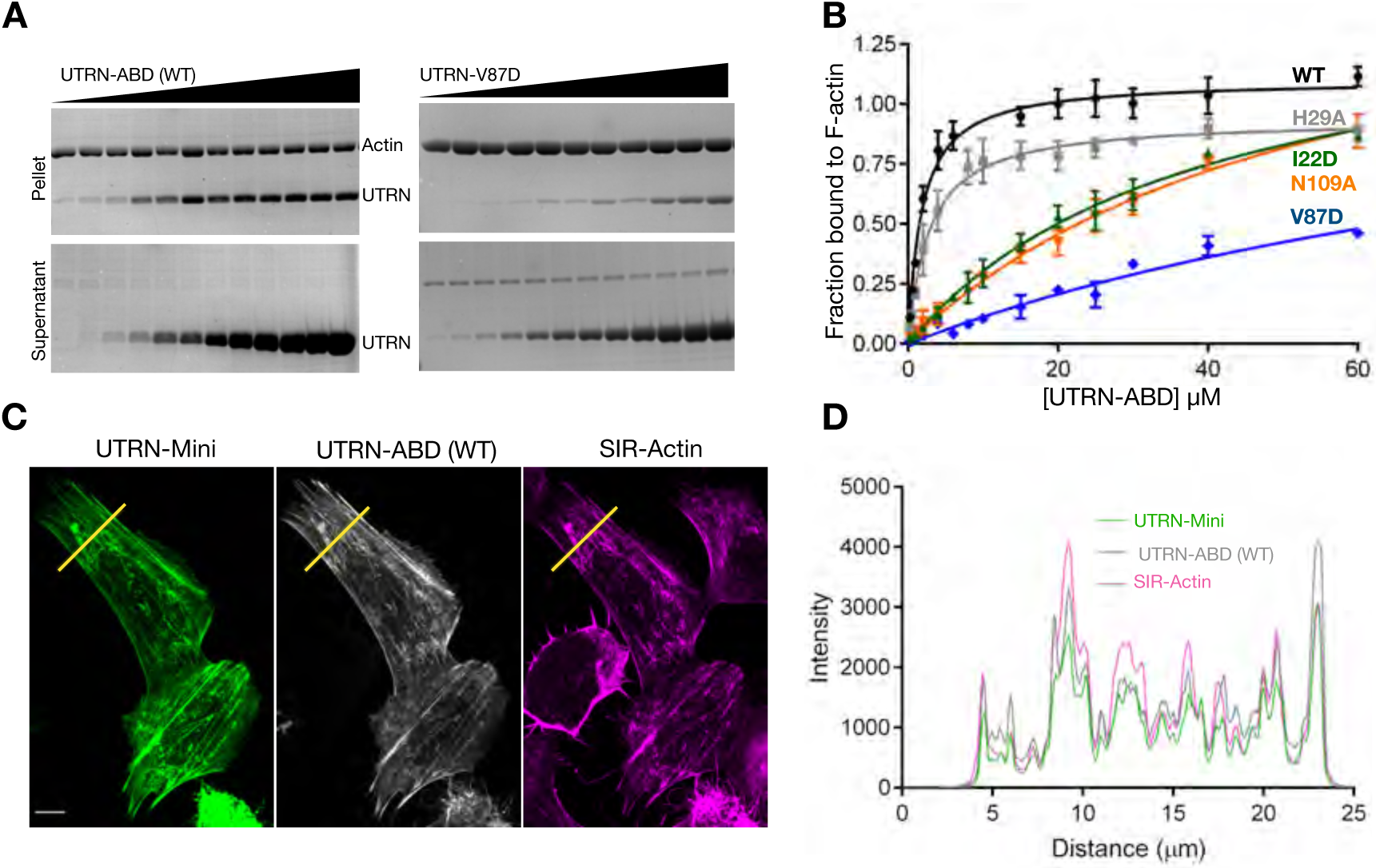
Mutation analysis of utrophin:F-actin binding interface. **A.** Representative Coomassie stained SDS-PAGE gels of UTRN-ABD and UTRN-V87D co-sedimentation with F-actin. Pellet (top) and supernatant (bottom) fractions of individual co-sedimentation reactions of increasing utrophin concentrations, uncropped gel images of all the co-sedimentation reactions are presented in Supplement Figure 6. **B.** Apparent K_d_ indicated was calculated from the titration data of co-sedimentation assays of utrophin wildtype (1.8μM) and mutants; H29A (2.8μM), I22D (38μM), V87D (>100μM) and N109A (55μM) as indicated. Data points for each concentration were averaged from 3 independent experiments, error bars represent S.D between independent experiments. **C.** Confocal images of U20S cells transfected with GFP tagged UTRN-mini and mCherry UTRN-ABD, stained with SIR-Actin shows F-actin structures, mainly stress fibers **D.** Co-localization analysis by intensity plot of GFP UTRN-mini, mCherry UTRN-ABD and SiR-Actin fluorescence using line scan of the region as indicated by yellow line in C.

Our mutagenesis study and structural model also suggests that ABD2’ and ABD2 sites of UTRN-ABD could be sufficient for F-actin interaction (Figure 4B, C & 5B). Therefore, we generated a truncated version called UTRN-mini encompassing amino acids 35-136, which was then tagged with GFP and compared with UTRN-ABD mcherry in U20S cell (Methods). Co-localization analysis shows that the UTRN-mini versus UTRN-ABD labelling of actin is nearly identical (Figure 5C & D). Together, from our structure and cell labeling studies we conclude that the CH1 domain of utrophin encompassing ABD2 and ABD2’ is sufficient for F-actin interaction.

## Discussion

Using cryoEM we have determined near atomic resolution structures of phalloidin-, lifeAct- and utrophin-bound to F-actin structures (Table 1). Representing the first high-resolution structural models of these most widely used F-actin cellular markers. Similar to most of the known F-actin structural models (3,4,14,15,30), the markers studied here do not induce any deviations in helical parameters of the actin filament (Table 1). By comparing the apo and phalloidin bound F-actin ADP structures, we conclusively show that phalloidin does not induce any conformation changes and closely resembles the ADP state of actin (Figure 1). This is in stark contrast to jasplakinolide, which shares the same binding site as phalloidin but causes the D-loop of actin monomers to adopt an open conformation, mimicking the ADP-Pi state(4). A recent structural study, which determined phalloidin-bound F-actin in different nucleotide states arrived at a similar conclusion (31).

Previous studies with utrophin tandem CH1 and CH2 domains, including an X-ray crystal structure(26), low-resolution electron microscopy models (23–25) and truncation studies (32) portray an ambiguous picture of utrophin:F-actin interaction and the actin binding sites. In our cryoEM reconstructions, we observe densities corresponding to the CH1 domain (amino acids 18-135) and an additional amino-terminal helix (amino acids 18-30), which was disordered in an earlier crystal structure (26) (Figure 4A and Supplement Figure 4). The utrophin CH1 domain architecture and the actin interaction regions are similar to the recently reported mutant FLNaCH1 (filamin) cryoEM structure (15). However, unlike the FLNaABD, where the CH2 domain showed weaker interaction and poor density map (15), we could not visualize utrophin CH2 domain in our reconstructions. Our structural observations of the CH1 domain are consistent with the biophysical characterization of UTRN-ABD, where upon actin binding of the CH1 and CH2 domain gets separated and adopts an open conformation (33,34), which is in contrast to the filamin tandem CH domains. The utrophin-F-actin structure also describes the important actin binding sites, for example the ABD2’ and ABD1, a helix unique to utrophin CH1 domain. Truncation studies guided by our structural model suggests that amino acids 35-136, i.e., the CH1 domain encompassing ABD2’ and ABD2 sites (UTRN-mini), could be sufficient for F-actin interaction. The UTRN-mini is a 101 amino acid protein that labels F-actin structures in cells (Figure 5C and D), which will occupy lesser footprint on F-actin compared to UTRN-ABD and could be advantageous in actin labelling experiments. Our UTRN-mini is in line with biochemical studies of utrophin CH1 domain (28) and filamin truncation studies, where FLNaCH1 shows similar actin labeling as FLNaABD (15). Utrophin, is also commonly used in biophysical experiments as a load in myosin motility assays (35), the mutations described here will be valuable to biophysicists in fine tuning the load exerted by utrophin in the motor assays. Together, we conclude that the CH1 domain of utrophin is an important element for F-actin interaction and can be used to label F-actin structures in cells (Figure 5C and D).

The lifeAct:F-actin complex cryoEM structure reveals that the lifeAct peptide adopts a 3-turn alpha-helix, as suggested by secondary structure prediction algorithms. The lifeAct interaction with actin is predominantly through hydrophobic contacts, encompassing two neighboring actin monomers. A key feature of this interaction is the overlapping site with D-loop (Figure 2 and 3), from which we hypothesized that lifeAct could sense the closed D-loop conformation, a hallmark of the F-actin ADP state. Our *in vitro* reconstitution experiments using phalloidin and jasplakinolide recapitulates the structural hypothesis that lifeAct detects F-actin in its closed D-loop state. Although our structural and biochemical studies support the importance of the D-loop in F-actin binding, lifeAct can interact with G-actin even more tightly (16), which is devoid of the D-loop from the adjacent actin monomer. Thus, a different binding state must exist for G-actin. We predict that in the absence of the D-loop, the charged carboxy terminus of lifeAct might play a dominant role in mediating additional electrostatic interactions with G-actin. However, in the case of the open D-loop state, the disruption of the hydrophobic pocket could sufficiently destabilize the F-actin interaction.

Previous cell biology experiments have indicated two limitations of lifeAct; 1. Affecting actin polymerization dynamics (17,20), and 2. Inability to label certain actin structures (8,19). The structural model provided here should help guide the creation of new lifeAct variants that will mitigate G-actin binding, further improving the suitability of the lifeAct probe. For the second limitation, the most common explanation is that lifeAct might be overlapping with other actin binding proteins. From our lifeAct work, we also suggest that the actin structures devoid of lifeAct signal might also reflect different biochemical or nucleotide states of actin, in this case specifically the ADP-Pi actin states.

With the advent of cryoEM, several actin binding proteins complexed with F-actin have been characterized (14,15,30,36). A common theme emerging from these structures is that SD1 and SD2 encompassing the D-loop region of actin is a preferred site for actin binding proteins (Figure 6). Our lifeAct and utrophin bound F-actin structures also show that they overlap with myosin, cofilin and coronin binding sites (30,37), but not with that of tropomyosin (Figure 6). Because the D-loop is the sole element that undergoes conformational changes in actin, several reports have proposed that actin binding proteins might sense the D-loop state. So far coronin and cofilin are known to sense the D-loop in open and closed conformation respectively (4,38). However, upon sensing the closed D-loop conformation, cofilin distorts the F-actin structure resulting in severing of the actin filament (37). Our work here describes the D-loop conformation sensing by lifeAct, which we also equate as a sensor for the ADP state of actin. Therefore, representing lifeAct as the first *bona fide* sensor for the closed D-loop and/or ADP nucleotide state of actin.

**Figure 6:**
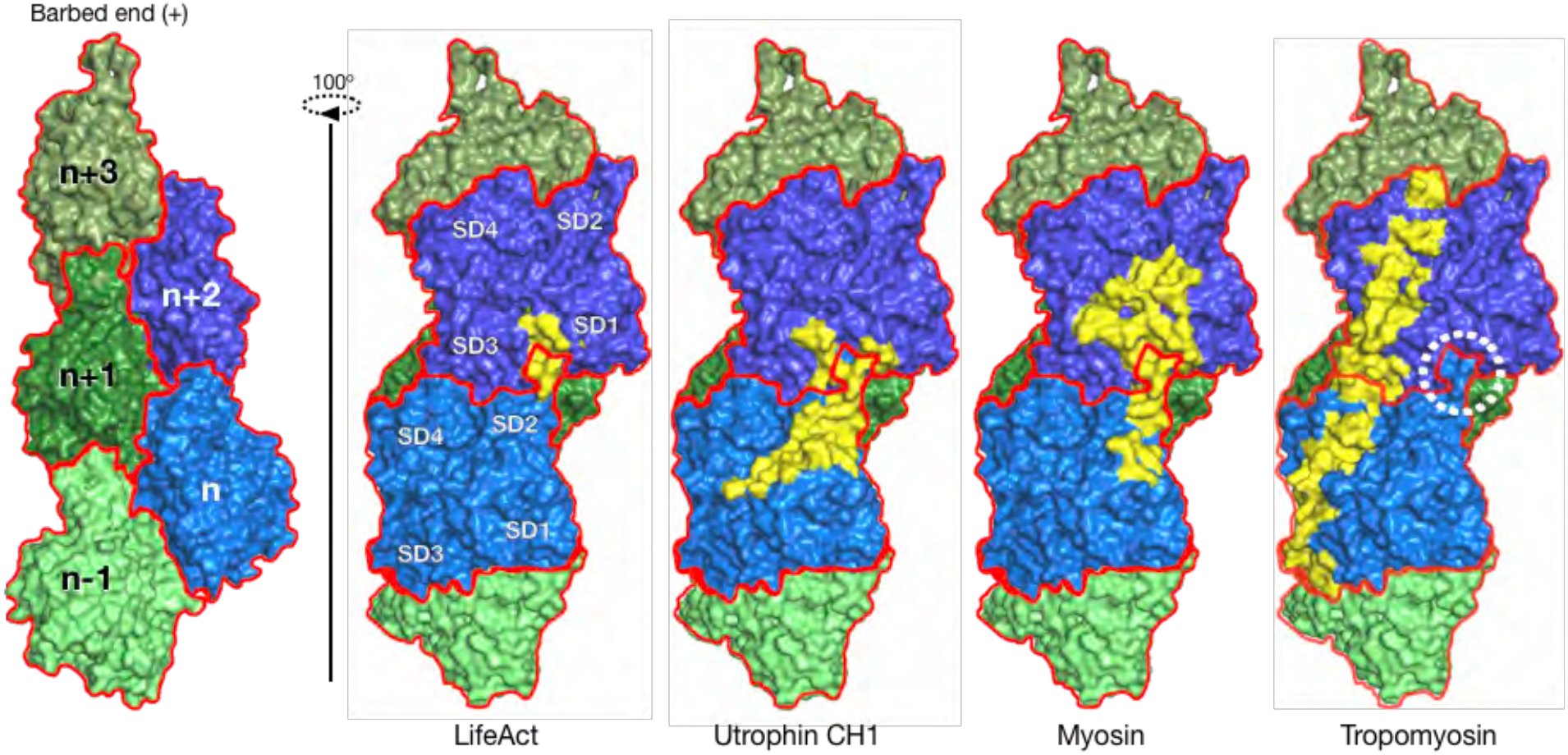
Interface comparison of UTRN, LifeAct with myosin and tropomyosin. Surface representation of F-actin with five monomers as marked. Footprint (in yellow) of actin monomers and respective residues interacting with lifeAct, utrophin CH1, myosin and tropomyosin as indicated and the D-loop is shown circled. The myosin and tropomyosin footprints were derived from PDB IDs 5JLH, 6C1D and 5JLF respectively.

In summary, our structural work combined with previous cell biological investigations of various actin markers offers insights into the nature of actin cell markers and their interactions with F-actin, providing an invaluable resource to the actin cytoskeleton community in choosing appropriate actin markers in their investigations.

## Materials and Methods

### DNA constructs and reagents

Human UTRN-ABD (amino acids 1-261) was cloned in pET28a vector with amino-terminal His tag, using GFP-UtrCH (addgene plasmid #26737) as a template. UTRN-ABD mutations were generated in the same vector by Quickchange site directed mutagenesis (Strategene). mcherry-UTRN-ABD and eGFP-UTRN-mini (amino acids 35-136) was cloned in mcherry and eGFP pCMV vector respectively. pLenti-LifeAct-EGFP BlastR was a gift from Ghassan Mouneimne (Addgene plasmid #84383). LifeAct mutations were created in the pLenti-LifeAct-EGFP BlastR using methodology as mentioned for UTRN. Alexafluor-568-phalloidin (Thermofischer Scientific Cat. No. A12380) and SiR-actin (Spriochrome Cat. No. Cy-SC001) were purchased. FAM-LifeAct peptide was custom synthesized from LifeTein, USA.

### Protein purification

6×His-tagged UTRN-ABD and mutants were expressed in *Escherichia coli* Rosetta DE3 strain and induced with 0.25mM IPTG overnight at 200C. Bacterial cells were pelleted and resuspended in lysis buffer (50mM Tris-Cl pH-7.5, 150mM NaCl, 20mM Imidazole, 0.1 % Tween-20 and Protease inhibitor cocktail tablet (Roche, Cat. No. 04693159001). The cells were lysed using sonication and the lysate was clarified at 18000 rpm for 30mins. The supernatant fraction containing proteins were loaded on 5ml His-Trap column (GE Healthcare) and eluted with a linear gradient of elution buffer containing 40mM to 500mM Imidazole, 500mM NaCl, 5mM beta mercaptoethanol. The utrophin protein fractions were pooled, concentrated and loaded on to the Superdex-200 16/600 column, preequilibrated with 50mM Tris-HCl pH-7.5, 150mM NaCl, 2mM TCEP, 0.1 % Tween-20. Pure fractions were concentrated using 3kDa MWCO centrifugal filter unit (Millipore), flash frozen in liquid nitrogen and store at −80°C until use.

### Actin co-sedimentation assays

Actin, purified from chicken breast (*Gallus gallus*) into G buffer (2 mM Tris pH 8, 0.2 mM ATP, 2 mM DTT, 0.2mM CaCl2) using Spudich lab protocol (39). G-actin was polymerized in F-actin buffer (25 mM Tris-Cl, 200 mM KCl, 2 mM MgCl2, 1 mM ATP) for 2 hrs at room temperature. Polymerized actin (7.5&μM) was titrated with increasing amounts of UTRN-ABD and mutants in co-sedimentation assay buffer (10mM Tris-Cl pH 8, 0.5 mM ATP, 0.2 mM DTT,2 mM MgCl2, 50 mM KCl). The mixture was incubated at room temperature for 20 mins, and spun at 100,000g for 30 mins in a Beckman TLA-100 rotor. Supernatants were collected, and protein pellets were suspended in equi-volume of co-sedimentation assay buffer. The supernatant, pellet and input samples were loaded in a 10% SDS–PAGE gel for separation. Gels were stained with Coomassie blue and scanned using the iBright FL1000 (Invitrogen). Fiji ImageJ was used for densitometric analysis. Data points were fitted to a one-site binding model using Prism software (GraphPad) to calculate the apparent binding affinity and stoichiometry as described earlier (28).

### Sample and grid preparation for cryo-EM

Freshly prepared G-actin was used for polymerization. For F-actin-phalloidin complex, actin was polymerized in F-actin buffer (10mM Imidazole pH 7.4, 200mM KCl, 2mM MgCl2, 1mMATP) at room temperature for 2 hrs and then Alexafluor-568-phalloidin (1/2 ratio) was mixed and incubated overnight at 4°C. For F-actin-utrophin and F-actin-lifeAct complex, polymerization was induced with KMEI buffer (50mM KCl, 1mM MgCl2, 0.5mM ATP, 1mM EGTA,10mM Imidazole pH 7.5) at 4°C overnight.

In the case of F-actin–phalloidin, we applied 3.0-3.5μl of sample onto a freshly glow-discharged Quantifoil Au 1.2/1.3, 300 mesh grids. Grids were prepared inside the Vitrobot. All grids were incubated for 30–60 s at >95% humidity, then blotted for 3-3.5 s. Immediately after blotting, the grids were plunge-frozen in liquid ethane. For F-actin-utrophin, 5-8 μM of F-actin was applied on to Au 1.2/1.3 grid and then 4-5 molar excess of utrophin was mixed to it, after 30-60s wait, it was blotted for 3.0-3.5 s and plunge-frozen in liquid ethane. For F-actin-lifeAct, Au 0.6/1.0 grid was used and same amount of F-actin with excess molar concentration of lifeAct peptide was used for sample preparation.

### Cryo-EM data collection

The datasets were collected on a Cs-corrected FEI Titan Krios G3 transmission electron microscope equipped with an FEG at 300 kV with the automated data-collection software EPU (ThermoFisher Scientific) at the National CryoEM facility, Bangalore. Images of the F-actin-phalloidin, F-actin-UTRN-ABD and F-actin-LifeAct were collected with a Falcon III detector operated in linear mode at a nominal magnification of 59,000X and a calibrated pixel size of 1.38 Å. In all cases, we acquired one images per grid hole. Table 1 contains the details on exposure time, frame number and electron dose for all the data sets.

### Data processing and model building

Unaligned frame images were manually inspected and evaluated for ice and filament quality. After manual removal of bad images, the remaining movie micrographs were motion corrected with either by Unblur (40) or by MotionCor2 inbuilt in Relion 3.0 (41). CTF estimation was performed with GCTF (Zhang K, 2016) on the full-dose motion-corrected sums. For all the datasets, filaments were manually selected and processed with Relion 3.0 (41). We used a box size of 353Å for F-actin-ADP apo, phalloidin bound and F-actin-Utrophin datasets and 320Å for the F-actin-lifeAct with the interbox distance of 29Å. After segment extraction, we performed 2D classification in Relion to remove bad segments. To further remove partially decorated filament, we did helical 3D classification using F-actin (EMDB-1990) as a reference, which was low-pass filtered to 30-35Å, to avoid reference bias. The best decorated 3D classes were combined and used for refinement using same reference as above with the sampling rate of 1.8°. All refinement steps were performed with soft mask containing 75-80% of the filament. To further improve the resolution, we performed CTF refinement and Bayesian polishing of lifeAct and utrophin bound F-actin data sets (42). The polished particles were refined for obtaining the final map.

We used F-actin structure (PDB-6BNO) as a starting atomic model in Chimera(43) to fit the F-actin model in the map for all data sets. Phalloidin coordinates were used from PDB-6D8C (15). For F-actin-utrophin complex, utrophin CH1 domain was taken from crystal structure PBD – 1QAG (26). Coot (44) was used for model building for all data sets and real space refined using Phenix (45). All structural models were validated in MolProbity (46) and PDBe site.

### Cell imaging

Wild type U2OS cells were obtained as a gift from Prof. Satyajit Mayor’s lab, NCBS Bangalore, India. For all the experiments, U2OS cells were cultured in McCoy’s 5A (Sigma Aldrich, M4892) media supplemented with 2.2g/l sodium bicarbonate, 10% fetal bovine serum (FBS) and 1x PenStrep (cat. no. 15-140-122 Gibco Fisher Scientific) in a humidified 37°C incubator with 5% carbon di oxide. Around 20,000 to 30,000 cells were seeded in ibidi glass bottom dishes (Cat. No. 81218, Ibidi) and transfection was carried out at 60-70% confluency of the cells with total of 1ug of plasmid DNA used for transfection. All the transfection experiments were carried out with jet prime transfection reagent (cat. no. 114-15 polypus transfection) as described in manufacture’s protocol. 500ng plasmid of UTRN-mini construct (UTRN 35-136) tagged with N-terminus EGFP were co-transfected with 500ng plasmid of mcherry tagged UTRN-ABD in the 10% serum containing media. The transfection media was changed with fresh media after 4-6 hours of transfection. Cells were imaged post 24hr of transfection with SiR-actin (Cat. no. CY-SC001 Spirochrome kit, Cytoskeleton, Inc) staining in the complete media. All the images were obtained at 60× oil objective (1.42NA) with 2048*2048 frame size and 0.5-1μm optical sections on FV3000 Olympus confocal microscope equipped with 488, 561 and 640 laser for GFP, cy3 and cy5 channels, respectively. All the images obtained through Olympus software and were analyzed on Fiji ImageJ.

### *In vitro* actin labeling assay and TIRF microscopy

Flow chambers of ~10ul volume were prepared using double-sticky tape, coverslips and cover glass. The flow chamber was incubated with Protein G (Sigma, Cat. No. 08062) for 10 minutes followed by anti-his antibody (Sigma, Cat. No. 11922416001) for another 10 mins. After washing with KMEI buffer without ATP, UTRN-ABD-6×His was flowed to attach actin filament with the coverslip. The Phalloidin-Actin-568 and SiR-actin 640 F-actin were prepared separately and added together with different concentration of FAM-lifeAct. The mixture was incubated in tube for 5-10 minutes and flowed in the chamber for visualization. Flow chambers were imaged at 100× oil objective 1.49NA under the total internal reflection mode using Nikon Ti2 H-TIRF system with 488, 561 and 640 laser lines. The images were acquired for all the three channels near glass surface sequentially with appropriate spectrum filter sets using s-CMOS camera (Hammamatsu Orca Flash 4.0) controlled by NIS-elements software. All images and data were analyzed using Fiji ImageJ software.

## Acknowledgements

The authors wish to thank the Jim Spudich and members of the Sirajuddin lab for comments on manuscript. The authors acknowledge the National CryoEM Facility at Bangalore Life Science Cluster and funding by B-life grant from Department of Biotechnology and the Central Imaging and Flow Facility (CIFF) at NCBS/inStem/CCAMP campus. A.K is supported by DBT-Research Associate Fellowship (No. 40796915460). This work was funded by NCBS core grants and SERB-Ramanujan Fellowship to VKR. inStem core grants from Department of Biotechnology, India, CEFIPRA (5703-1), Wellcome Trust-DBT India Alliance Intermediate Fellow (IA/I/14/2/501533) and EMBO Young Investigator award to MS.

## Author Contributions

AK and MS conceived the project. AK, MGJ and VKR performed cryoEM work and biochemical analysis. AK, SK and MS performed cell biology and TIRF experiments. VKR and MS supervised the project. AK and MS wrote the paper and all authors commented on the manuscript.

## Data Availability

CryoEM maps and coordinates are deposited in EMDB and PDB under following code; PDB XXXX and EMD-XXXX for F-actin:ADP apo, PDB XXXX and EMD-XXXX for F-actin:ADP phalloidin, PDB XXXX and EMD-XXXX for F-actin:lifeAct and PDB XXXX and EMD-XXXX for F-actin:utrophin. The datasets for this study are available from the corresponding author upon reasonable request.

---

**Supplement Figure 1:**
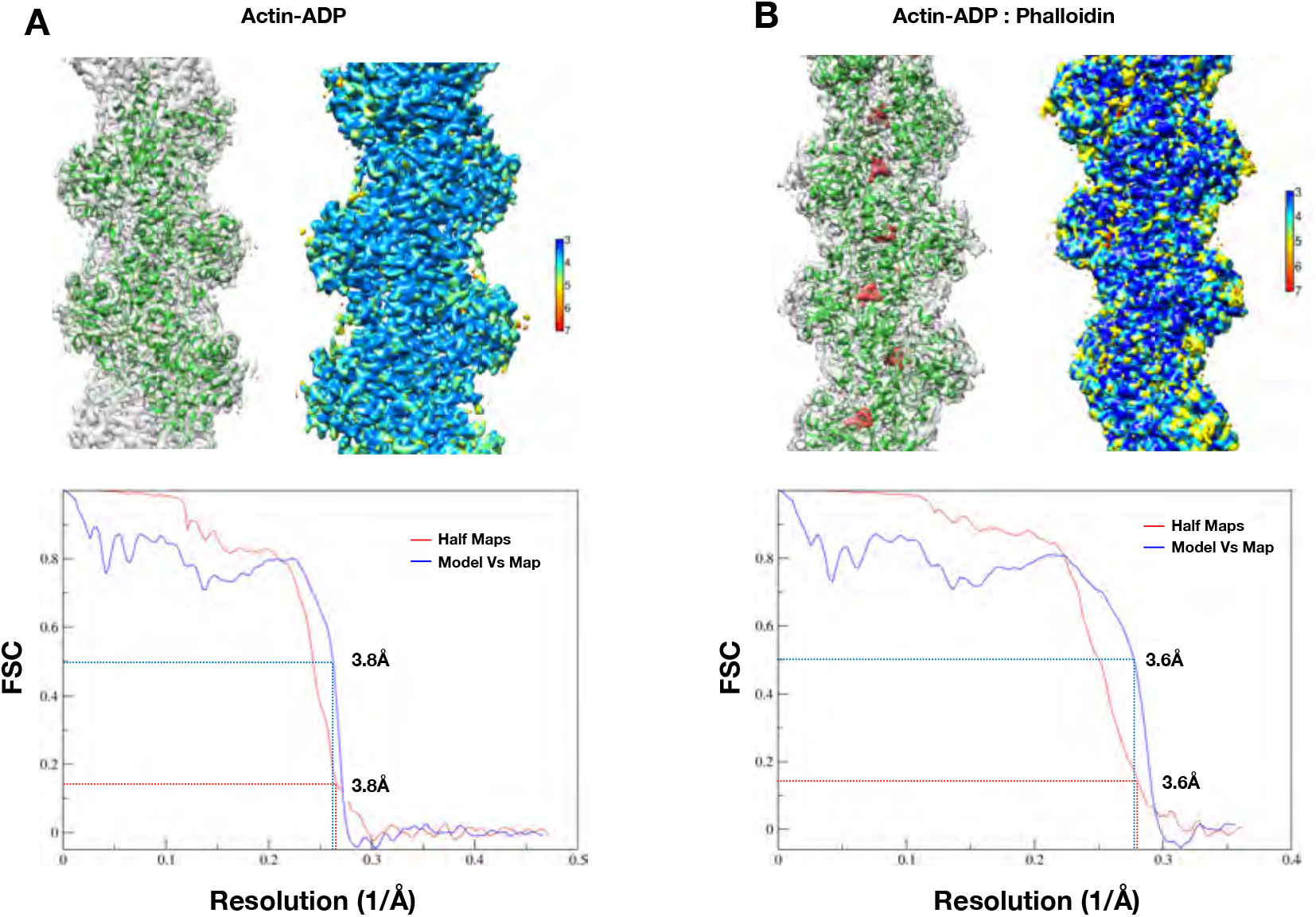
CryoEM structure validation and quality control. **A.** and **B.** CryoEM structure and local resolution map of actin:ADP and actin:ADP-phalloidin structures. Overlay of the model in the EM map versus map and the local resolution of maps (from Resmap) for actin:ADP and actin:ADP-phalloidin respectively with color gradient chart **C. and D.** FSC plots of the half-maps and the map versus model for the Actin-ADP and Actin-ADP-phalloidin respectively.

**Supplement Figure 2:**
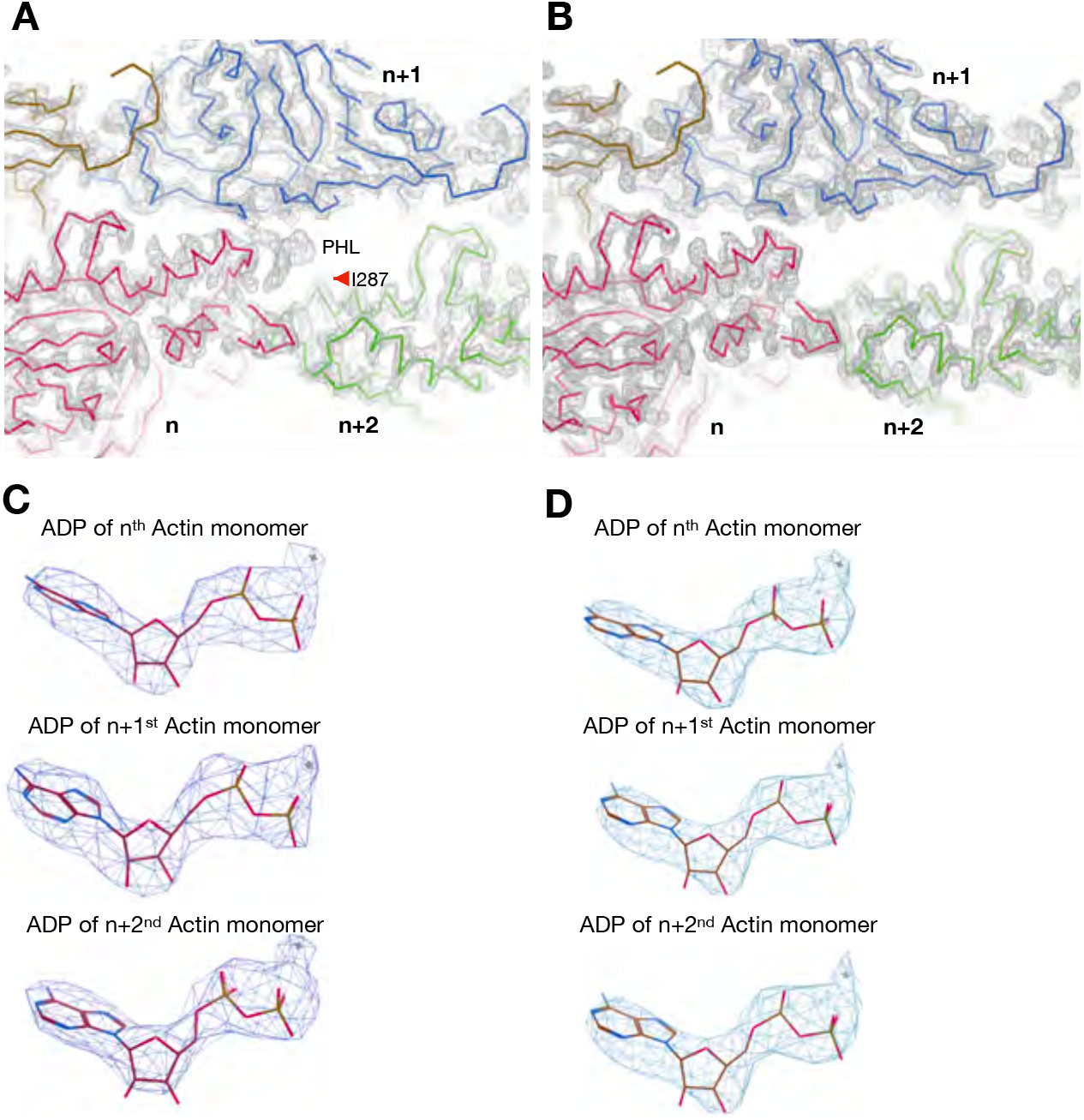
F-Actin ADP, apo versus phalloidin bound maps. **A.** and **B.** F-actin ADP apo versus phalloidin maps. The density of phalloidin is indicated with PHL and the I287 residue is marked as red arrow. **C** and **D**. ADP of respective structures of actin monomers as indicated.

**Supplement Figure 3:**
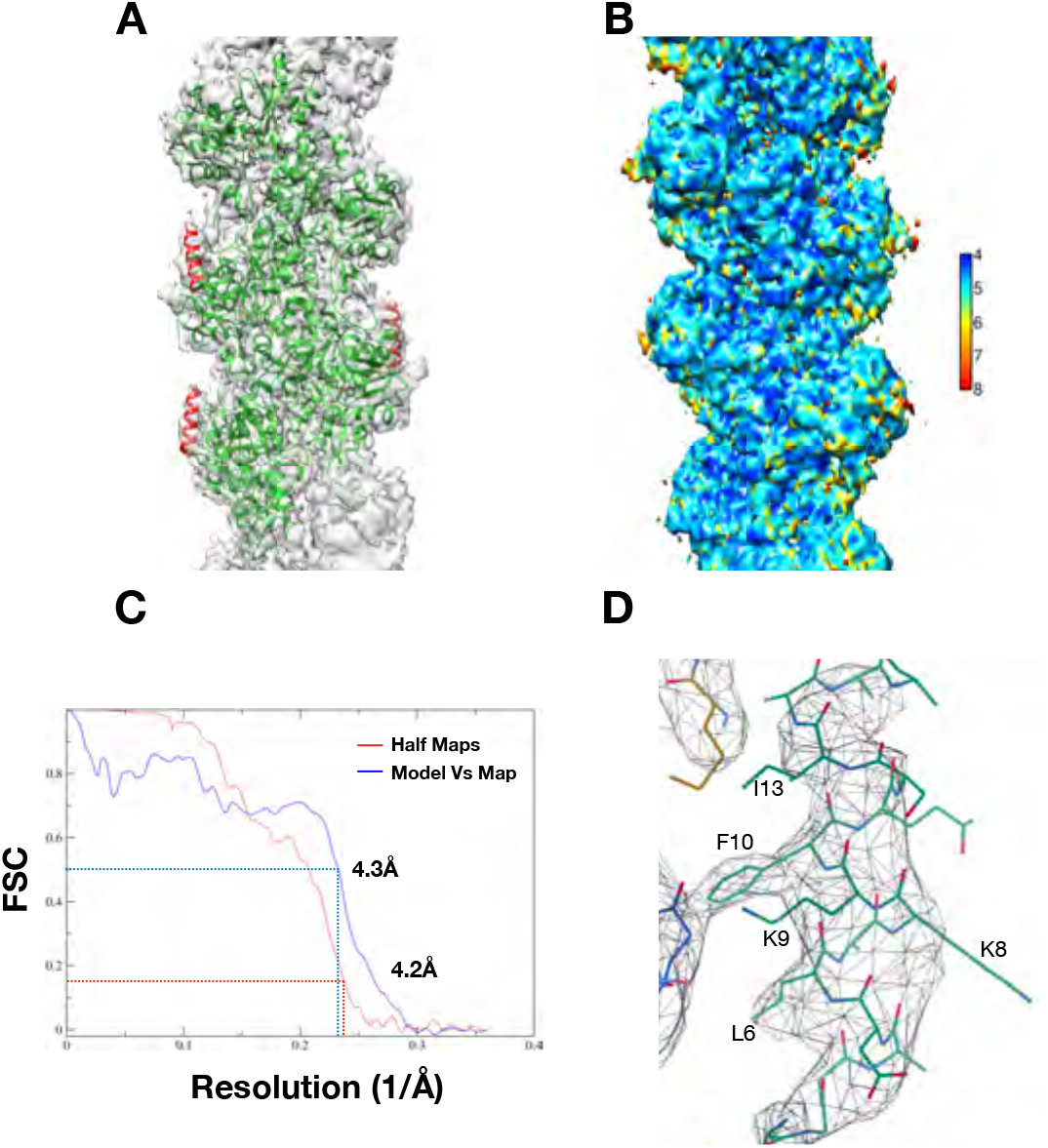
LifeAct maps with F-actin binding site. **A.** Overlay of model with map, actin and lifeAct are colored as green and red respectively. **B.** Local resolution of lifeAct bound F-actin map with color gradient chart determined with Resmap. **C.** FSC plots of map versus model for F-actin:lifeAct structure. **D.** Closer view of lifeAct map with key residues marked.

**Supplement Figure 4:**
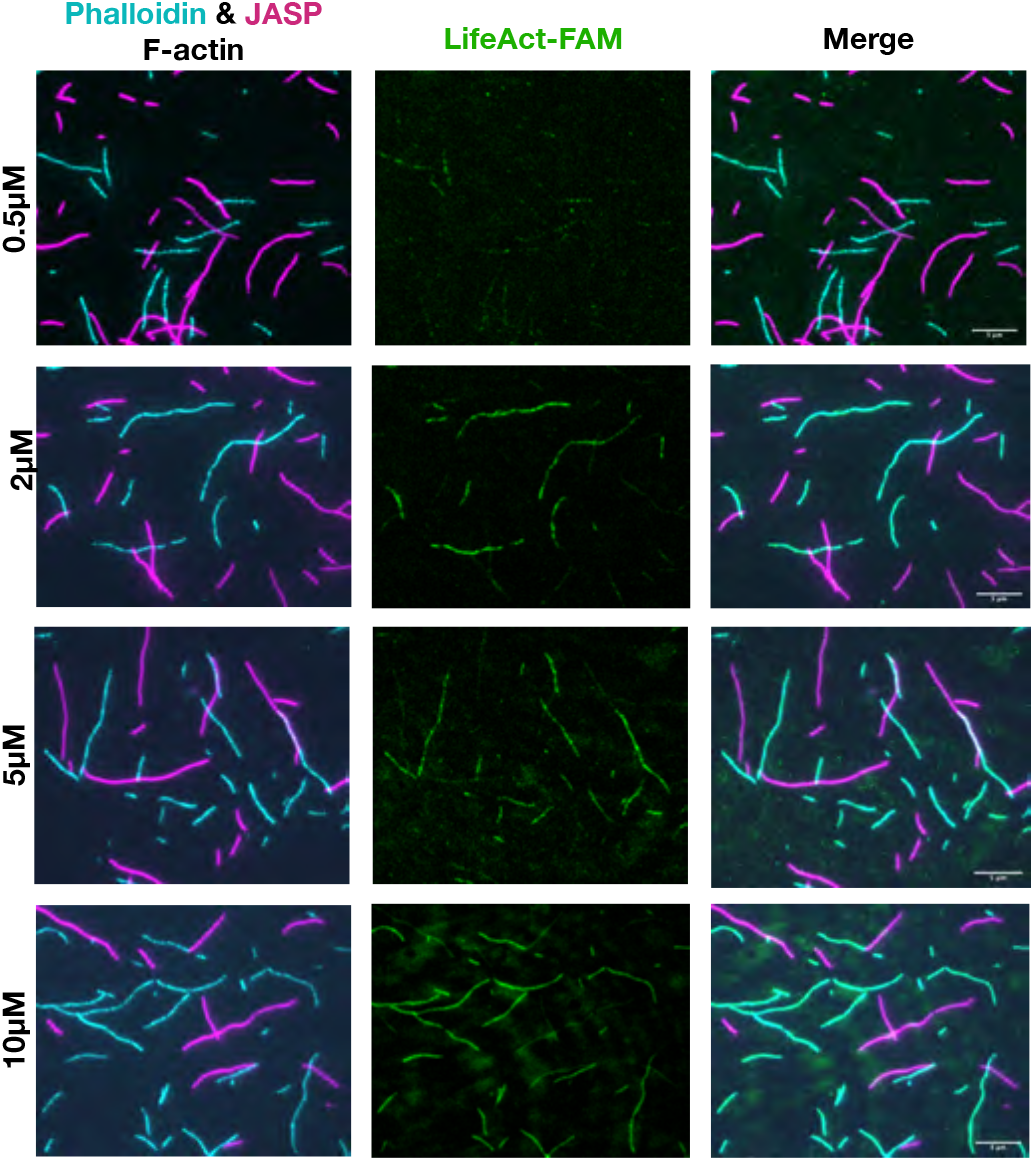
LifeAct TIRF data supplement material. Representative TIRF images of lifeAct versus phalloidin and jasplakinolide bound F-actin for lifeAct concentrations as indicated.

**Supplement Figure 5:**
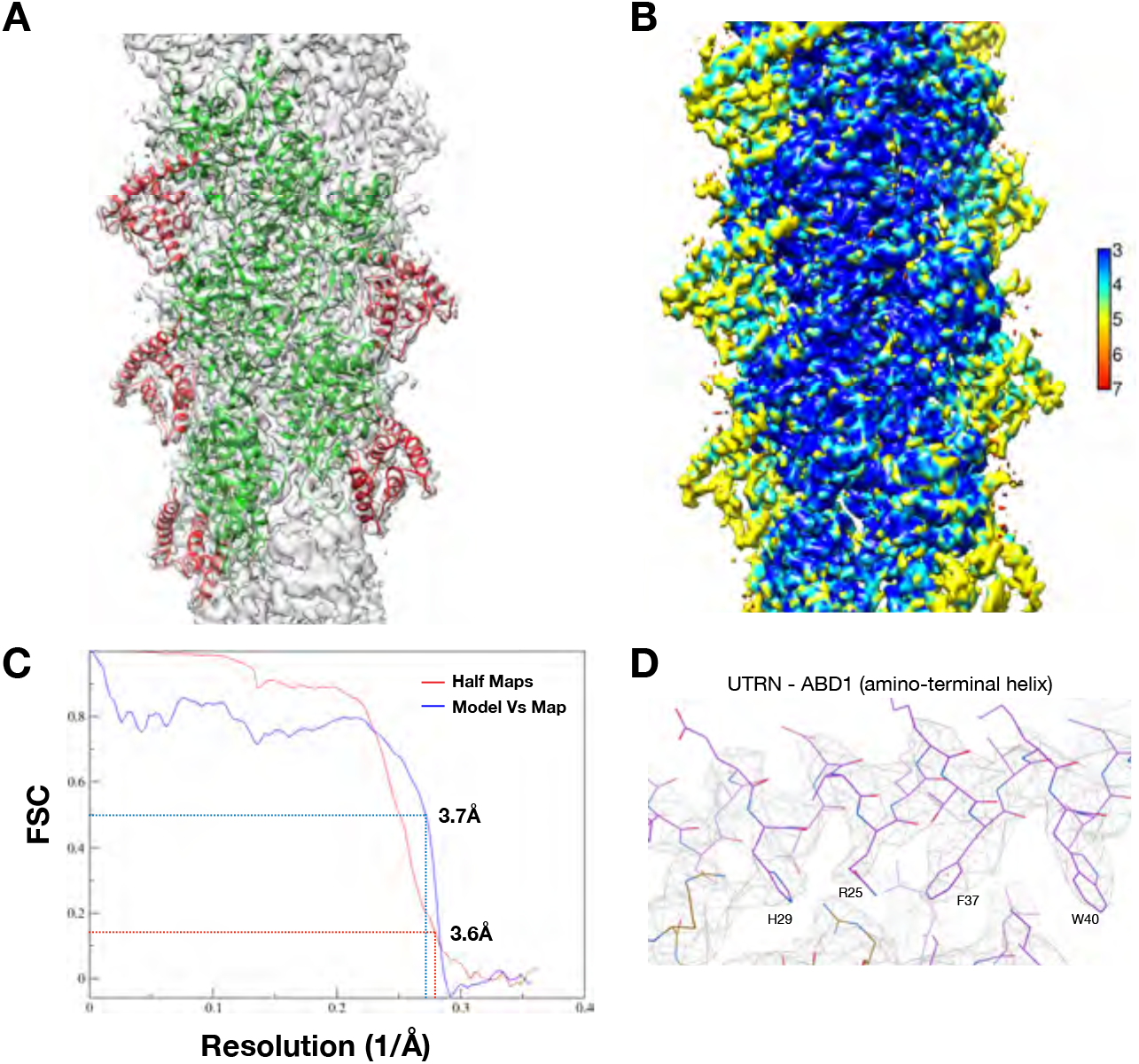
Utrophin bound F-actin map. **A.** Overlay of model with map, actin and utrophin CH1 are colored as green and red respectively. **B.** Local resolution of utrophin CH1 bound F-actin map with color gradient chart as indicated. **C.** FSC plots of the half-maps and the map versus model for F-actin:utrophin CH1 structure. **D.** Closer view of ABD1 (amino-terminal helix) utrophin map with key residues marked.

**Supplement Figure 6:**
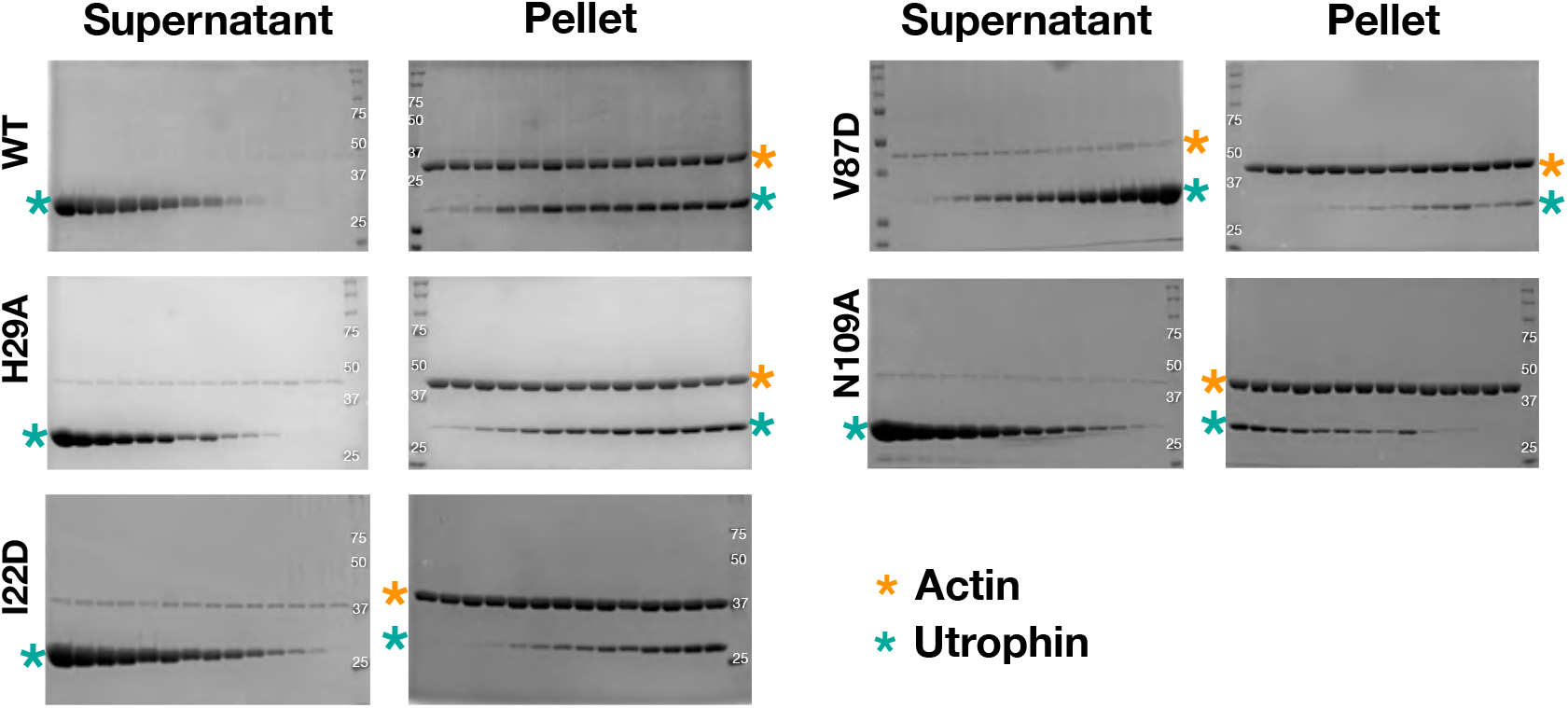
UTRN mutation co-sedimentation gels. Uncropped gel images of co-sedimentation experiments with F-actin versus utrophin wildtype and mutants used in calculating the K_d_ shown in Figure 5B.

